# C4BP occludes the non-opsonic interaction of *Neisseria gonorrhoeae* with human neutrophil CEACAMs

**DOI:** 10.64898/2026.01.19.700458

**Authors:** Mary W. Broden, Jutamas Shaughnessy, Frida Mohlin, Amaris J. Cardenas, Anna M. Blom, Sanjay Ram, Alison K. Criss

## Abstract

*Neisseria gonorrhoeae* (Gc) causes the sexually transmitted infection gonorrhea, an urgent public health concern. Gc infection elicits a robust neutrophil response and serum leakage, but Gc has developed specialized defenses to evade both complement and neutrophils. We recently reported that the classical complement pathway inhibitor C4b-binding protein (C4BP) binds to Gc and reduces phagocytic killing by neutrophils in a complement-independent manner. Here, we used a Chinese hamster ovary (CHO) expression system and engineered C4BP constructs to define the underlying molecular mechanisms. C4BP inhibited interactions between opacity protein (Opa)-expressing Gc and carcinoembryonic antigen-related cell adhesion molecules (CEACAMs), receptors that drive non-opsonic phagocytosis of Gc by neutrophils. The degree of C4BP-mediated inhibition varied among CEACAMs. By using wild-type and chimeric CEACAMs, we found C4BP was more inhibitory towards the granulocyte-restricted CEACAM3 than the ubiquitously expressed CEACAM1, which we ascribed to CEACAM3’s shorter extracellular domain. C4BP also inhibited the association between Opa-expressing Gc and the GPI-anchored CEACAM6. Molecules containing C4BP domains 1 and 2 fused to IgM (C4BP-IgM) or to a hexameric IgG Fc construct (C4BP-Hexa-IgG), proteins similar in diameter and degree of multimerization to native C4BP, inhibited the association of Opa-expressing Gc with CEACAM3-CHO cells to the same degree as C4BP, while C4BP domains 1 and 2 fused to dimeric Fc (C4BP-IgG) did not. C4BP-IgM, but not C4BP-IgG bearing mutations to abrogate Fc gamma receptor interactions, blocked Opa-mediated phagocytosis by primary human neutrophils. These results support a model in which C4BP occludes Opa-CEACAM interactions, which protects Gc from phagocytic killing by neutrophils.

## Introduction

*Neisseria gonorrhoeae* (gonococcus; Gc) is a Gram-negative, obligate human pathogen that causes nearly 87 million new cases of the sexually transmitted infection gonorrhea annually (Rowley et al., 2019). Sharply increasing resistance to all classes of antibiotics, in conjunction with the lack of a licensed vaccine or protective immunity from previous infection, makes the understanding of Gc pathogenesis to inform new therapeutics a pressing need (Unemo et al., 2019). Symptomatic Gc infection characteristically results in a robust influx of neutrophils to infected sites (Evans, 1977; Ovcinnikov & Delektorskij, 1971; Stevens & Criss, 2018). Neutrophil transmigration and release of inflammatory effectors result in epithelial damage, followed by serum leakage to infected sites (Stevens & Criss, 2018). To survive in this highly inflammatory environment, Gc has developed human-specific defenses against both soluble and cellular components of the innate immune system [reviewed in (Lewis & Ram, 2020; Palmer & Criss, 2018)].

One such defense is the recruitment of human C4b-binding protein (C4BP) to the bacterial surface via outer membrane protein porin (PorB) (Ram et al., 2001). C4BP inhibits the classical complement pathway, which protects Gc from antibody-mediated serum bactericidal activity (Ram et al., 2001). C4BP is a large (570 kDa) protein whose most common isoform circulating in serum consists of 7 alpha chains and 1 beta chain, each of which is composed of CCP (complement control protein) repeats (Dahlback et al., 1983). The most distal (or N-terminal) CCP repeat (CCP1) of each alpha chain contains the binding site for the Gc porin PorB (Jarva et al., 2007; Ram et al., 2001), which accounts for 60% of outer membrane protein content (Johnston et al., 1976). Our lab recently reported that C4BP has an additional complement-independent role in Gc pathogenesis: C4BP binding to Gc protects Gc from non-opsonic phagocytosis and killing by neutrophils (Werner et al., 2023). We showed that C4BP is both necessary and sufficient for human serum-mediated protection of Gc from neutrophils, and C4BP binding to Gc blunts the Opa-induced oxidative burst of neutrophils (Werner et al., 2023).

The major driver of non-opsonic interaction of Gc with neutrophils is the binding of gonococcal opacity-associated (Opa) family proteins to neutrophil carcinoembryonic antigen-related cell adhesion molecules (CEACAMs), a family of Ig-like receptors. CEACAMs 1, 3, and 6 are expressed by neutrophils and are capable of interaction with various Opa proteins (Chen & Gotschlich, 1996; Chen et al., 1997; Gray-Owen, Dehio, et al., 1997; Gray-Owen, Lorenzen, et al., 1997; Popp et al., 1999; Virji, Makepeace, et al., 1996; Virji, Watt, et al., 1996). CEACAMs 1 and 3 are transmembrane proteins containing intracellular signaling motifs, while CEACAM6 binds the cell membrane via a glycosylphosphatidylinositol (GPI) anchor. Each CEACAM contains one or more extracellular Ig-like domains. Opa binding is mediated by the CEACAM N-terminal Ig-like domain, termed IgV (Ig-variable) (Virji, Watt, et al., 1996).

CEACAM1 is a broadly expressed, intracellular tyrosine-based inhibitory motif (ITIM)-containing receptor whose engagement on immune cells generally elicits Src homology domain-containing protein tyrosine phosphatase (SHP)-dependent inhibitory signaling (Chen et al., 2008; Lee et al., 2008; Thomas et al., 2023). The full-length version of human CEACAM1 has 3 Ig constant-like (IgC) domains between the transmembrane region and the N-terminal IgV domain. CEACAM3 is exclusively expressed by the granulocytes of primates and has been described as a decoy receptor that may have evolved for phagocytic host defense; while pathogens exploit the binding to CEACAM on epithelial cells for colonization, they are engulfed and killed when interacting with CEACAM3 on neutrophils (Adrian et al., 2019). CEACAM3 is an intracellular tyrosine-based activating motif (ITAM)-containing phagocytic receptor. CEACAM3 engagement by Opa induces receptor clustering and ITAM phosphorylation (McCaw et al., 2003). Phosphorylated ITAM recruits spleen tyrosine kinase (Syk), which signals through downstream effectors to induce phagocytosis, ROS production, and cytokine release (McCaw et al., 2004; Sarantis & Gray-Owen, 2007; Schmitter et al., 2004; Sintsova et al., 2014). CEACAM6 is GPI-anchored and does not contain a cytoplasmic signaling domain (Kolbinger et al., 1989), and its role in neutrophil responses to Opa+ Gc remains largely unexplored.

The mechanism by which C4BP binding to Gc inhibits Opa-mediated phagocytosis and downstream activation leading to ROS production in primary human neutrophils has not yet been defined. C4BP does not interact directly with Opa or CEACAM (Werner et al., 2023). However, C4BP is a large, multimeric protein complex that binds PorB, which is the most abundant protein in the Gc outer membrane (Johnston et al., 1976). Moreover, C4BP specifically prevented phagocytosis of CEACAM3-binding Opa+ Gc, but not IgG-opsonized, Opa-negative Gc, which instead engages Fc gamma receptors (Werner et al., 2023). Based on these observations, we hypothesized that C4BP sterically occludes Opa-CEACAM interactions at the Gc-neutrophil interface. In the present study, we used CEACAM-transfected cell lines, primary human neutrophils, and recombinantly generated C4BP-based constructs to assess how C4BP affects Opa-dependent interactions of Gc with neutrophil CEACAMs and to identify the underlying molecular determinants.

## Results

### CEACAM-CHO system for quantifying Opa-CEACAM interactions

To assess the role of C4BP in modulating interactions between individual Opa proteins and individual CEACAMs, we heterologously expressed human CEACAMs in pgsD-677 CHO cells. This CHO cell derivative is heparan sulfate proteoglycan (HSPG)-deficient (Lidholt et al., 1992) and lacks all known Opa binding partners as it is a cell line of hamster, not human origin. The cDNA for full-length CEACAM1 or CEACAM3 (Illustrated in Fig 1A), or an empty control vector, was introduced into these cells (called CHO cells hereafter) by stable transfection. Opa-mediated interaction of Gc with these cells was assessed using derivatives of strain FA1090 in which all *opa* genes were deleted (Opaless, designated here as “Opa-”), and into which single, constitutively expressed, non-phase-variable *opa* genes were reintroduced (Ball and Criss 2013). For this study, Gc constitutively producing strain FA1090 OpaD (Ball & Criss, 2013), strain FA1090 OpaI (Alcott et al., 2022), or strain MS11 Opa60 (Martin et al., 2016) in the Opaless background were used. These Opa proteins have been reported to bind both CEACAM3 and CEACAM1 (Alcott et al., 2022; Werner et al., 2020). For simplicity, the Gc used here will be referred to by the name of the Opa protein each produces (e.g. “OpaD”).

**Figure 1:**
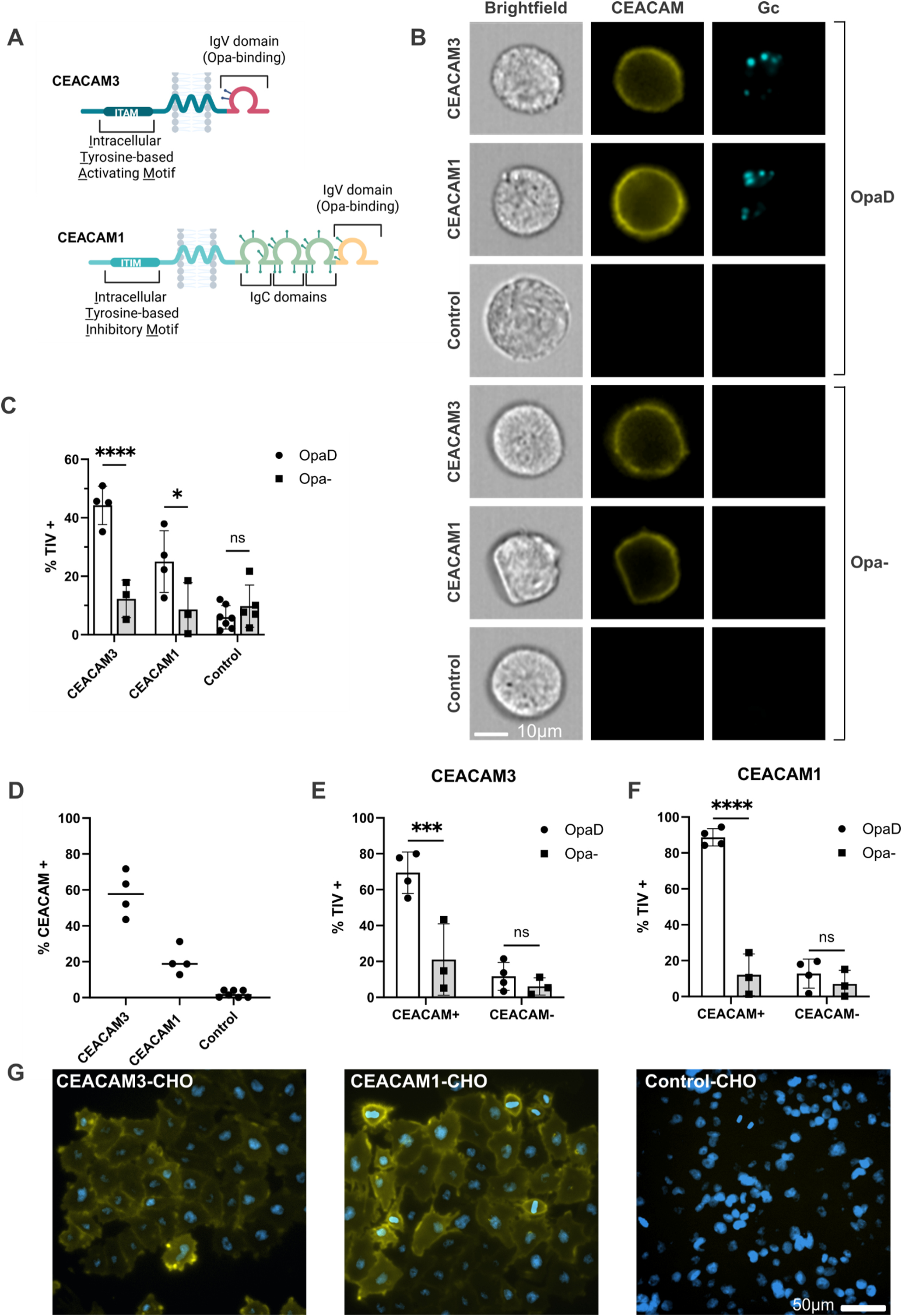
CEACAM-transfected Chinese hamster ovary cells as a platform for analyzing receptor-mediated interactions with Opa-expressing *N. gonorrhoeae*. (A) Schematic of CEACAM1 and CEACAM3 proteins used in this study. (B-F) Unsorted CEACAM1, CEACAM3, or empty vector transfected (Control)-CHO cells were infected with Tag-it-Violet^TM^ (TIV)-labeled Gc (OpaD or Opa-) at MOI 15. Cells were labeled with anti-Pan CEACAM antibody, and CEACAM expression and bacterial association with cells were assessed by imaging flow cytometry. (B) Representative imaging flow cytometry images of bacterial binding to CEACAM1, CEACAM3, or Control-CHO cells. Scale bar = 10 µm. (C) Bacterial association with unsorted CHO cells in each population, quantified by TIV fluorescence. (D) Percent of CEACAM-expressing cells in each cell population. Each dot indicates one biological replicate that reports the mean percent of cells that are CEACAM-expressing in that population; the median of these replicates is indicated by the line. (E-F) Bacterial association with CEACAM1-CHO or CEACAM3-CHO cells by TIV positivity when population is gated by CEACAM expression. In C, E, and F, bars depict the mean +/− SD for the indicated number of replicates. Groups were compared using two-way ANOVA followed by Tukey’s post-hoc test for statistical significance. * p<0.05, *** p<0.001, **** p<0.0001, ns = not significant. (G) CEACAM-positive cells were isolated from the CEACAM1-CHO and CEACAM3-CHO populations above by flow cytometric cell sorting. CEACAM expression in the sorted populations was confirmed by immunofluorescence microscopy (mouse anti-Pan CEACAM antibody followed by anti-mouse-AF555 secondary, DAPI counterstain). Scale bar = 50 µm.

To test if CEACAM-CHO cells can be used to quantify Opa-CEACAM interactions, confluent monolayers of CEACAM1, CEACAM3, or control CHO cells were infected with Tag-it-Violet^TM^ (TIV)-labeled Opa- or OpaD, and bacterial association was quantified by imaging flow cytometry (Fig 1B-F). As expected, OpaD associated with the CEACAM1- and CEACAM3-CHO cells significantly more than Opa-Gc (Fig 1C). Control cells not expressing CEACAM showed low levels of association with Gc, regardless of Opa expression (Fig S1). These results confirmed that in this model system, Opa and CEACAM expression are both required for Gc-cell interaction.

Cells in the bulk transfected populations did not homogenously express CEACAM on their surface (Fig 1D). When cells were gated based on CEACAM expression, OpaD associated significantly more with the CEACAM-positive population of CEACAM3-CHO cells and CEACAM1-CHO cells than the CEACAM-negative population, while Opa-had minimal association with the cells in either gate (Fig 1E-F), further confirming the specificity of Opa-CEACAM interactions in this system. To remove the CEACAM-negative background cells from the stably transfected populations, CEACAM-positive CHO cells were isolated by fluorescence-activated cell sorting (FACS). As shown in Fig 1G, the resulting populations were predominantly CEACAM-positive, and they were used for all subsequent experiments.

### C4BP more potently blocks the interaction of Opa+ *Neisseria gonorrhoeae* with CEACAM3 than with CEACAM1

To test how C4BP impacted the interaction of Opa+ Gc with CEACAMs, the bacterial inoculum was optimized using a spot counting algorithm that was applied to images generated from imaging flow cytometry. For each CEACAM-producing cell line and Opa-producing strain of Gc, the multiplicity of infection (MOI) was experimentally selected so that 40-60% of cells were Gc-positive after 45 min, e.g. contain 1 or more Gc. MOI for Opa-strains was matched to the OpaD MOI used for each cell line. Vector-only Control-CHO cells were infected at an MOI of 5. Opa- or OpaD was incubated with purified C4BP at 100 µg/mL, the minimum concentration that was experimentally determined to yield maximal binding to Gc (Fig. S3A), then washed prior to addition to the cells.

Incubation with C4BP abolished OpaD-CEACAM3 interactions, decreasing the level of bacterial association to that of Opa-Gc (Fig 2A). In contrast, C4BP reduced OpaD-CEACAM1 interactions to 72% of the total bacterial binding without C4BP (Fig 2B). The association of OpaD with Control-CHO cells remained at background levels regardless of C4BP (Fig S1A). To compare the magnitude of the effect of C4BP between conditions, these results were reported as the relative association with CHO cells, that is, the percent of Gc-positive cells infected with C4BP-bound bacteria, divided by the percent of cells positive for Gc without C4BP. A relative association value of 1 indicates that C4BP has no effect; the greater the effect of C4BP on Gc-CHO interaction, the lower the relative association (< 1). By this metric, comparing the relative effect of C4BP on Opa-CEACAM interactions, C4BP inhibited OpaD-CEACAM3 populations significantly more than it inhibited interactions with CEACAM1 (Fig 2C).

**Figure 2:**
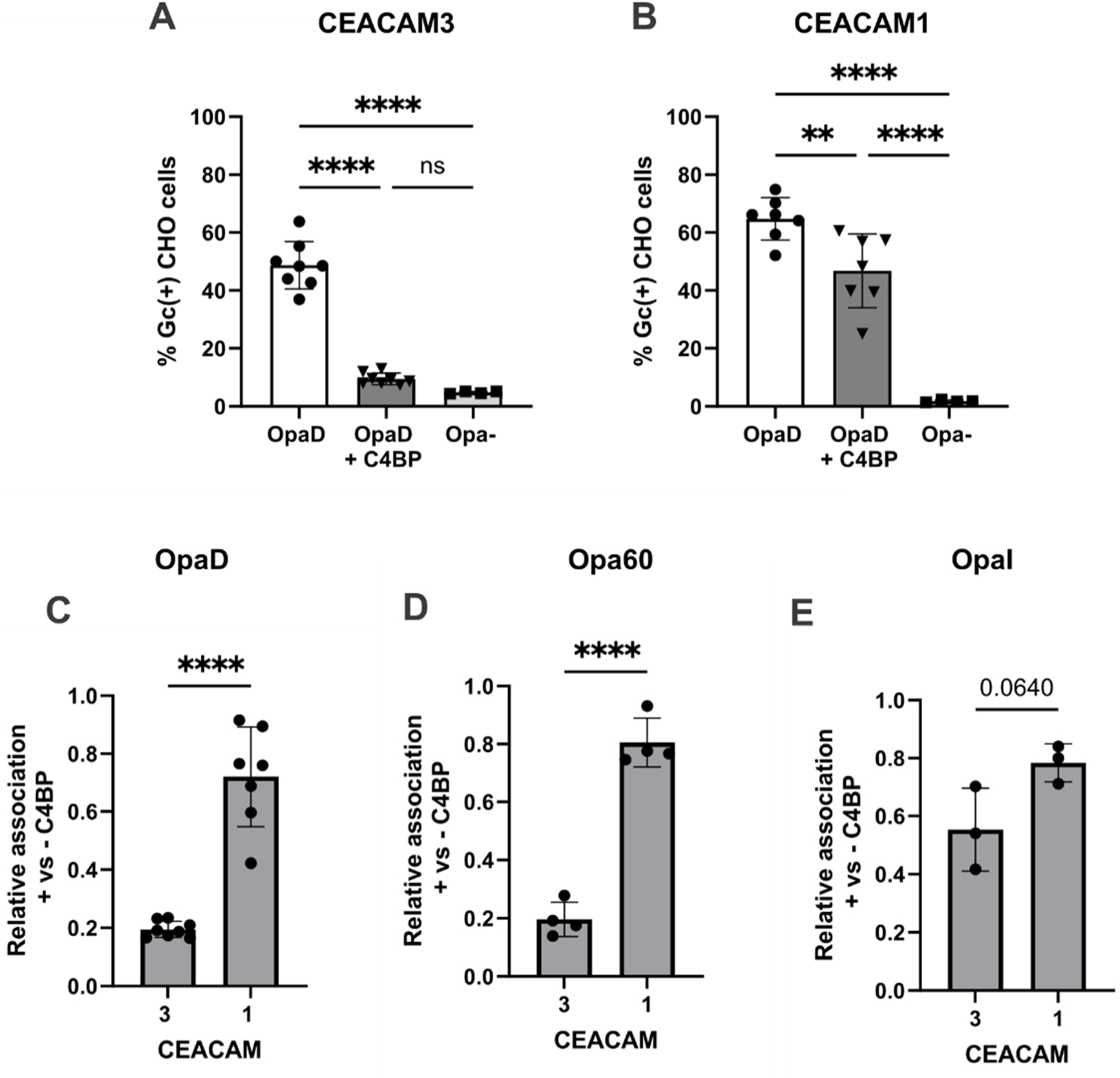
C4BP more potently inhibits interaction of Opa+ *N. gonorrhoeae* with CEACAM3 than with CEACAM1. CEACAM1 or CEACAM3-CHO cells were infected with TIV-labeled Gc. Cells were analyzed by imaging flow cytometry. The number of Gc per cell was quantified using a spot counting algorithm, and results are presented as the percent of Gc-positive CHO cells (e.g. containing at least 1 bacterium). A minimum of 3500 cells were analyzed. (A) CEACAM3-CHO cells were infected with OpaD, OpaD pre-incubated with 100 µg/mL C4BP (OpaD + C4BP), or Opa-Gc, each at MOI = 10. (B) CEACAM1-CHO cells were infected with OpaD ± C4BP or Opa-Gc, each at MOI = 5. (C-E) The effect of C4BP on the interaction of OpaD (C), Opa60 (D), and OpaI (E) with CEACAM1- and CEACAM-3 CHO cells, presented as relative association. Relative association is defined as the percent of Gc-positive CHO cells infected with C4BP-bound bacteria, divided by the percent of Gc-positive cells infected with bacteria without C4BP. Experiments with Opa60 in (D) used MOI = 10 for CEACAM3-CHO cells and MOI = 5 for CEACAM1-CHO cells. Experiments with OpaI in (E) used MOI = 5 for both CEACAM3-CHO cells and CEACAM1-CHO cells. Graphs depict the mean +/− SD. Statistical significance was determined in (A-B) by one-way ANOVA with Tukey’s multiple comparisons and in (C-E) by unpaired T-test. **p<0.01, ****p<0.0001, ns = not significant.

Similar to the findings with OpaD, C4BP significantly inhibited Opa60 binding to CEACAM3-CHO cells, with a statistically significant but less pronounced effect on association with CEACAM1-CHO cells (Fig 2D, S2A-B). Compared with OpaD and Opa60, C4BP had less of an effect on the association of OpaI with CEACAM-expressing cells (Fig 2E, S2C-D). However, C4BP still affected the association of OpaI with CEACAM3 more strongly than with CEACAM1. There was a minimal background level of association of all Gc strains with the control CHO cells (Fig S1).

From these experiments, we conclude that C4BP blocks Opa-CEACAM interactions, with the magnitude of inhibition being higher for CEACAM3 than CEACAM1.

### CEACAM structural investigations

Based on the structural differences between CEACAMs 1 and 3 (Fig 1A), we evaluated three features of CEACAM3 that could explain how C4BP more strongly inhibited Gc association with it than with CEACAM1: 1) The ability of CEACAM3 to signal intracellularly; 2) The amino acid sequence of the Opa-binding N-terminal, IgV domain; 3) The extracellular length of the receptor. To this end, we generated CHO cells expressing selected CEACAM1 and 3 mutants and chimeras, and the effect of C4BP on Opa-dependent bacterial binding was compared with parental CEACAM3 and CEACAM1 cells. cDNA sequences and detailed descriptions of each construct are listed in Table S1.

First, we generated CHO cells expressing CEACAM3-YYFF, which contains the mutations Y230F and Y241F (Fig 3A). These point mutations in the phosphorylatable tyrosines within the ITAM render CEACAM3 signaling-deficient without compromising overall structure (Billker et al., 2002; McCaw et al., 2003). C4BP abolished interactions of OpaD with CEACAM3-YYFF, with association similar to Opa-bacteria (Fig 3B), and there was no difference in the degree of C4BP-mediated decrease between CEACAM3-YYFF and wild type CEACAM3 (Fig 3C). These data show that the intracellular signaling capacity of CEACAM3 does not contribute to its ability to be blocked by C4BP.

**Figure 3:**
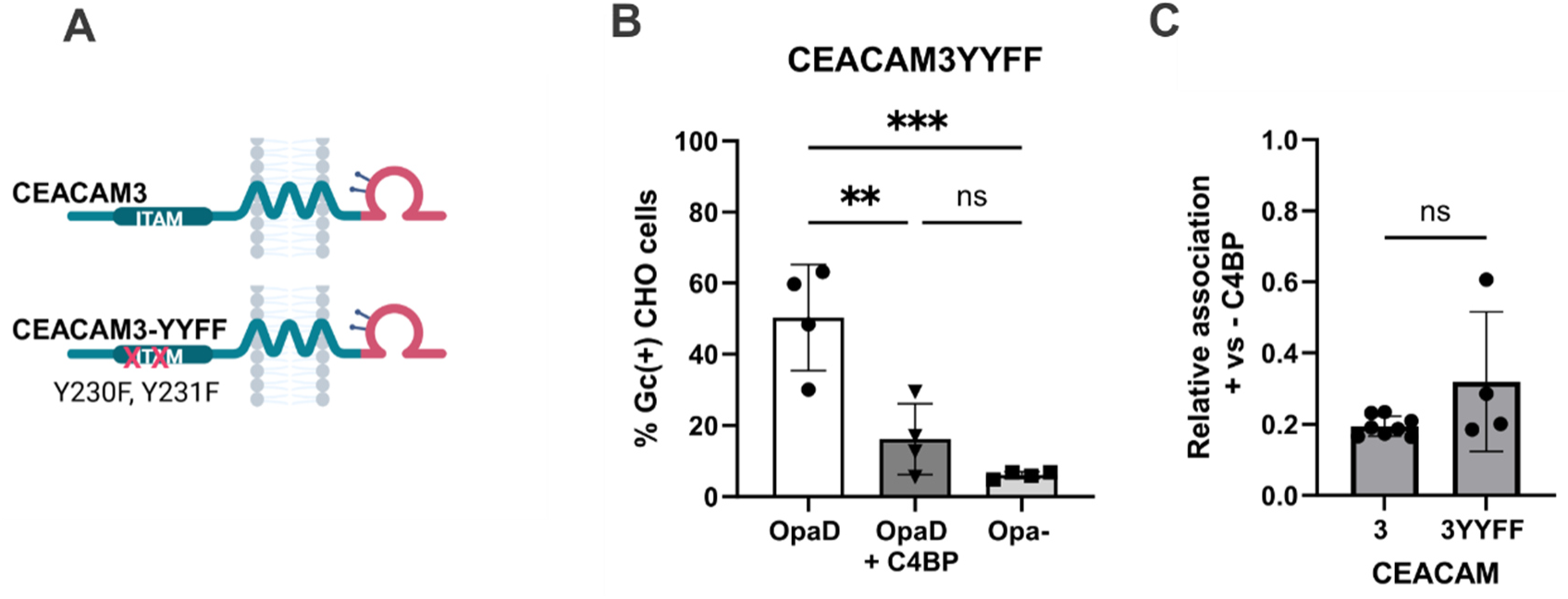
ITAM activity does not contribute to the inhibitory effect of C4BP on Opa-CEACAM3 interactions. (A) Schematic of full-length CEACAM3 and the CEACAM3-YYFF mutant, which lacks ITAM-dependent signaling ability. (B) CEACAM3-YYFF CHO cells were infected with OpaD ± C4BP or Opa- (MOI = 10 for each condition), and the percent Gc-positive cells was calculated by imaging flow cytometry as in Figure 2. (C) C4BP-dependent relative association of OpaD with CEACAM3-YYFF-CHO cells compared to CEACAM3-CHO cells (data from Fig 2C). Graphs depict the mean +/− SD. Statistical significance was determined in (B) by one-way ANOVA with Tukey’s multiple comparisons and by (C) unpaired T-test. ** p<0.01, *** p<0.001, ns = not significant.

To assess whether CEACAM3’s IgV domain sequence is responsible for the increased inhibitory effect of C4BP, we generated CHO cells expressing the chimeric CEACAM1-3N receptor, in which the N-terminal IgV domain of CEACAM1 is replaced by the IgV domain of CEACAM3 (Fig 4A). CEACAM1-3N interactions with OpaD were inhibited by C4BP (Fig 4B), to the same limited degree as measured for CEACAM1, and significantly less than measured for CEACAM3 (Fig 4C). We conclude that the IgV domain sequence does not affect how efficiently C4BP inhibits Opa-CEACAM interaction.

**Figure 4:**
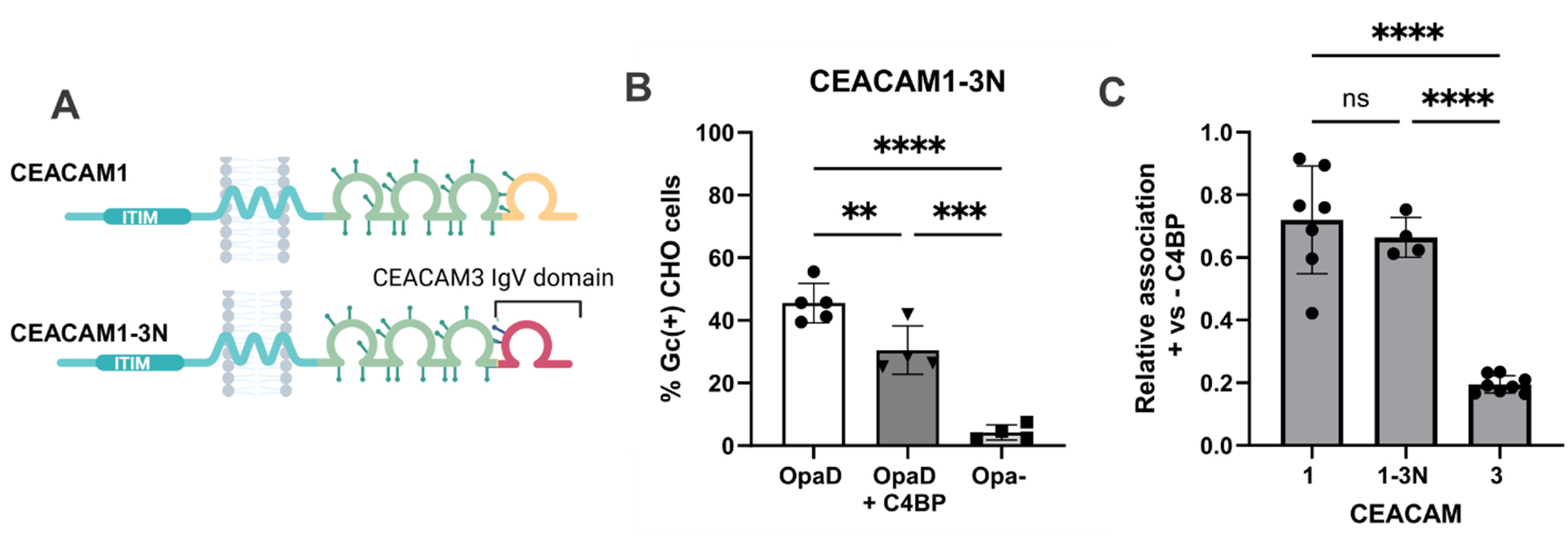
IgV domain sequence does not contribute to the inhibitory activity of C4BP on Opa-CEACAM interactions. (A) Schematic of the CEACAM 1-3N chimera relative to CEACAM1. (B) CEACAM1-3N-CHO cells were infected with OpaD ± C4BP or Opa- (MOI = 5 for each condition), and the percent of Gc-positive cells was calculated using imaging flow cytometry as in Figure 2. (C) C4BP-dependent relative association of OpaD with CEACAM1-3N-CHO cells compared to CEACAM1-CHO and CEACAM3-CHO cells (data from Fig 2C). Graphs depict mean +/− SD. Statistical significance determined by one-way ANOVA with Tukey’s multiple comparisons. ** p<0.01, *** p<0.001, **** p<0.0001, ns = not significant.

These results pointed to the length of the extracellular region of CEACAM as determining the magnitude of the inhibitory effect of C4BP on interactions of Opa with CEACAMs, which was tested in two ways. First, CHO cells were transfected to express chimeric CEACAM3 proteins in which the 3 IgC domains of CEACAM1 or 6 IgC domains of CEACAM5 were inserted after the N-terminal IgV domain. These constructs were named CEACAM3L and 3XL for long and extra-long, respectively (Fig 5A). The inhibitory effect of C4BP was less pronounced for CEACAM3L and 3XL than for parent CEACAM3 (Fig 5C-E); with increasing CEACAM3 length, there was a corresponding decrease in the inhibitory effect of C4BP. Conversely, CHO cells were generated that express a mutant of CEACAM1 with its 3 IgC regions deleted, named CEACAM1S (for “short” extracellular domain of CEACAM1) (Fig 5B). C4BP had a more prominent inhibitory effect on the interaction of OpaD with CEACAM1S-CHO cells compared with CEACAM1-CHO, although it did not fully recapitulate the level of inhibition measured with parental CEACAM3 (Fig 5F-G). Together, these results implicate the extracellular length of CEACAMs 1 and 3 in the magnitude of the inhibitory effect of C4BP on Opa-CEACAM interactions.

**Figure 5:**
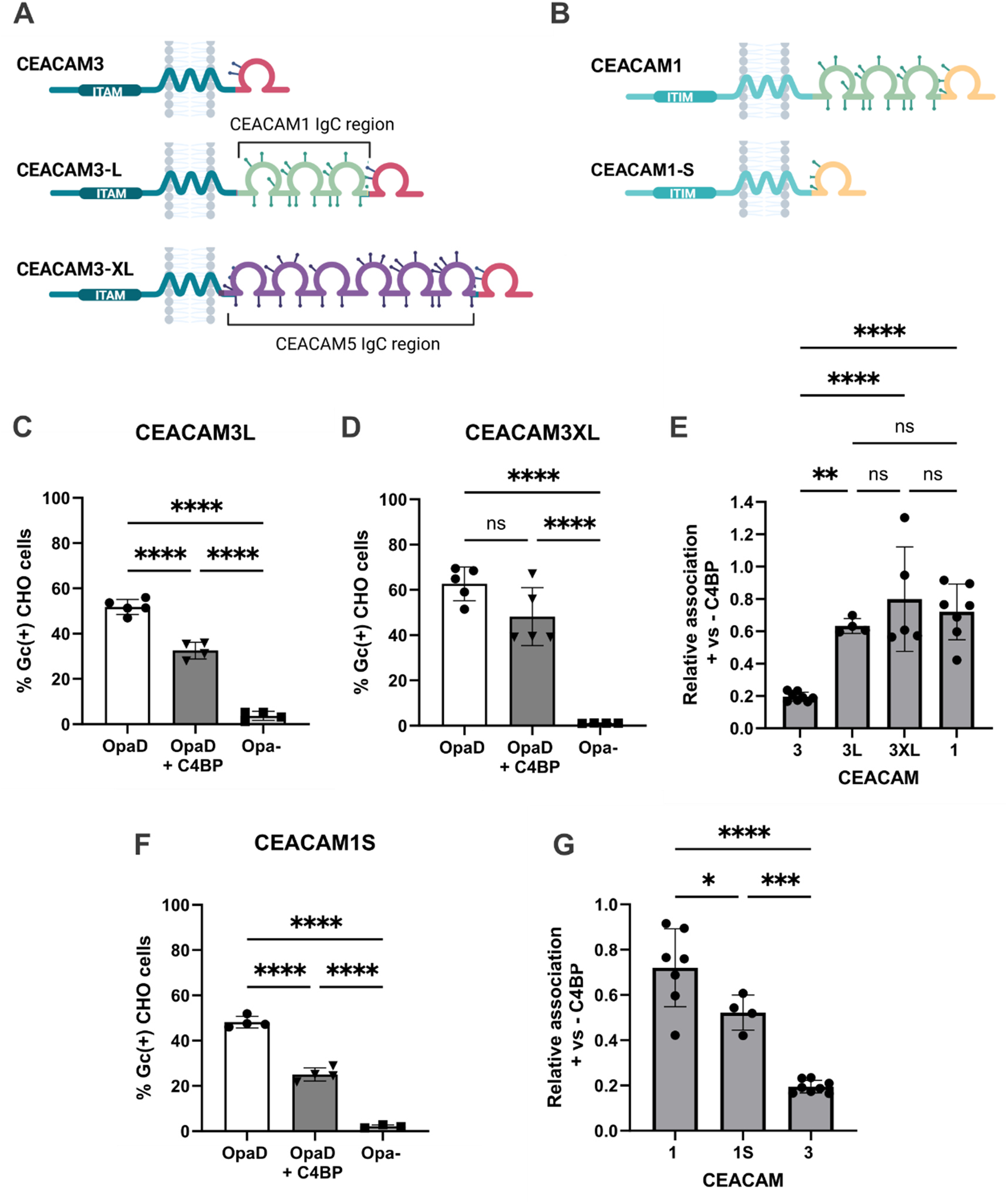
CEACAM extracellular length contributes to the inhibitory activity of C4BP on Opa-CEACAM interactions. (A) Schematics of CEACAM3L and CEACAM3XL relative to CEACAM3. (B) Schematic of CEACAM1S relative to CEACAM1. (C) CEACAM3L-CHO cells were infected with OpaD ± C4BP or Opa- (MOI = 5 for each condition), and the percent Gc-positive cells was calculated by imaging flow cytometry as in Figure 2. (D) CEACAM3XL-CHO cells were infected with OpaD ± C4BP or Opa- (MOI = 2.5 for each condition), and the percent Gc-positive cells was calculated using imaging flow cytometry as in Figure 2. (E) C4BP-dependent relative association of OpaD with CEACAM3L-CHO cells and CEACAM3XL-CHO cells compared to CEACAM3-CHO cells and CEACAM1-CHO cells (data from Fig 2C). (F) CEACAM1S-CHO cells were infected with OpaD ± C4BP or Opa- (MOI = 20 for each condition), and the percent Gc-positive cells was calculated using imaging flow cytometry as in Figure 2. (F) C4BP-dependent relative association of OpaD with CEACAM1S-CHO cells compared to CEACAM1-CHO and CEACAM3-CHO cells (data from Fig 2C). Graphs depict the mean +/− SD. Statistical significance was determined by one-way ANOVA with Tukey’s multiple comparisons. * p<0.05, ** p<0.01, *** p<0.001, **** p<0.0001, ns = not significant.

Neutrophils also express the Opa-binding CEACAM6, which is GPI-anchored rather than transmembrane (Fig 6A). Thus, we hypothesized that the effect of C4BP on the interaction of Opa+ Gc with CEACAM6 would differ from the effects on CEACAMs 1 and 3. Because OpaD does not bind CEACAM6 (Werner et al., 2020), we tested this hypothesis using the CEACAM6-binder Opa60 ((Werner et al., 2020), Fig 6). Addition of C4BP strongly inhibited the association of Opa60 with CEACAM6-CHO cells; in fact, inhibition was more potent than the effect on association with CEACAM3 or CEACAM1-CHO cells (Fig 6B-C). These data suggest that how CEACAMs are anchored to the membrane may also contribute to the extent of C4BP-mediated inhibition.

**Figure 6:**
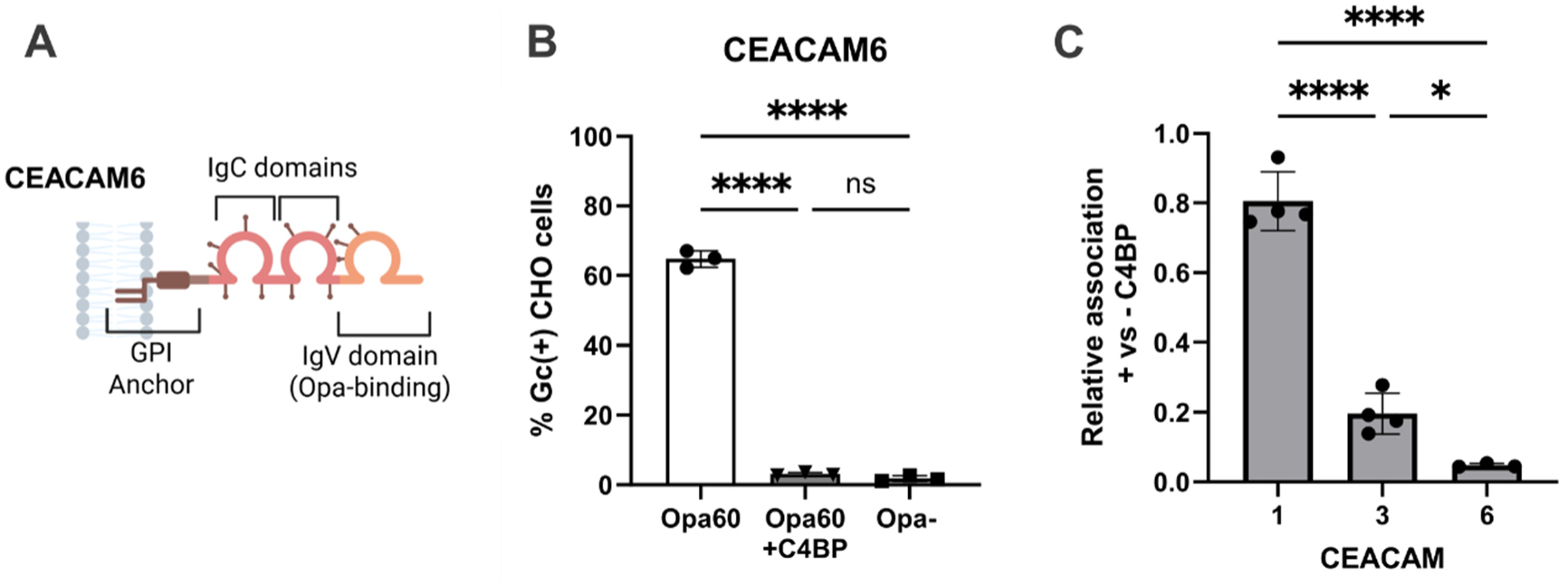
C4BP inhibits interactions of Opa+ *N. gonorhoeae* with CEACAM6. (A) Schematic of CEACAM6. (B) CEACAM6-CHO cells were infected with OpaD ± C4BP or Opa- (MOI = 5 for each condition), and the percent Gc-positive cells was calculated using imaging flow cytometry as in Figure 2. (C) C4BP-dependent relative association of Opa60 with CEACAM6-CHO cells compared to CEACAM1-CHO and CEACAM3-CHO cells (data from Fig 2D). Graphs depict mean +/− SD. Statistical significance determined by one-way ANOVA with Tukey’s multiple comparisons. * p<0.05, **** p<0.0001, ns = not significant.

### Size and degree of multimerization of C4BP contribute to the ability to block Opa-CEACAM3 interactions and neutrophil phagocytosis of Gc

To test how the size and degree of multimerization of C4BP impacts Opa-CEACAM3 interactions, we engineered an array of C4BP constructs. An alpha chain-only, recombinantly produced hexamer of C4BP inhibits non-opsonic phagocytosis of Opa+ Gc by neutrophils and is no different from purified C4BP at equimolar concentrations (Werner et al., 2023), allowing us to use alpha chain-based C4BP constructs to interrogate Opa-CEACAM interactions. These constructs retain CCP1, the distal C4BP alpha chain domain that is required for binding to gonococcal PorB, including the PorB1b allele found in strain FA1090 used here (Ram et al., 2001).

To test if replacing the core structure of C4BP with an unrelated large, branching structure would be sufficient to retain blocking ability, we employed two fusion proteins: C4BP-IgM and C4BP-Hexa-IgG. C4BP-IgM is a fusion protein built on a multimeric IgM Fc backbone, where the IgM Fab regions are replaced by CCP1-2 of human C4BP (Bettoni et al., 2019). C4BP-Hexa-IgG is built on an IgG1 backbone engineered to hexamerize by introduction of the C-terminal IgM tailpiece, and where its Fab domains are also replaced by CCP1-2 (Fig 7A). Binding of C4BP-IgM and C4BP-Hexa-IgG to Gc was confirmed using antibodies recognizing the Fc regions of the constructs (Fig 7B, S3B-C). Both C4BP-IgM and C4BP-Hexa-IgG blocked the interaction of OpaD with CEACAM3-CHO cells in a statistically significant manner, similar to native C4BP (Fig 7C). We then used a recombinant C4BP-IgG Fc fusion protein with monomeric Fc. C4BP-IgG is an IgG3-based construct where the Fab region is replaced with two copies of CCP1 (Fig 7A). As this construct does not contain the IgM multimerization sequence, it retains the smaller, dimeric structure of native IgG3. C4BP-IgG bound to Gc (Figs 7B, S3D), but unlike its multimeric counterparts, C4BP-IgG did not impede interactions between OpaD and CEACAM3-CHO cells (Fig 7C). Thus, the ability of native C4BP to block Opa-CEACAM interaction can be recapitulated by introducing the Gc-binding regions of C4BP on a similarly sized, unrelated backbone, but not on a smaller dimeric construct. Together, these results suggest that the size and degree of multimerization of C4BP contribute to its ability to block Opa-CEACAM3 interactions.

**Figure 7:**
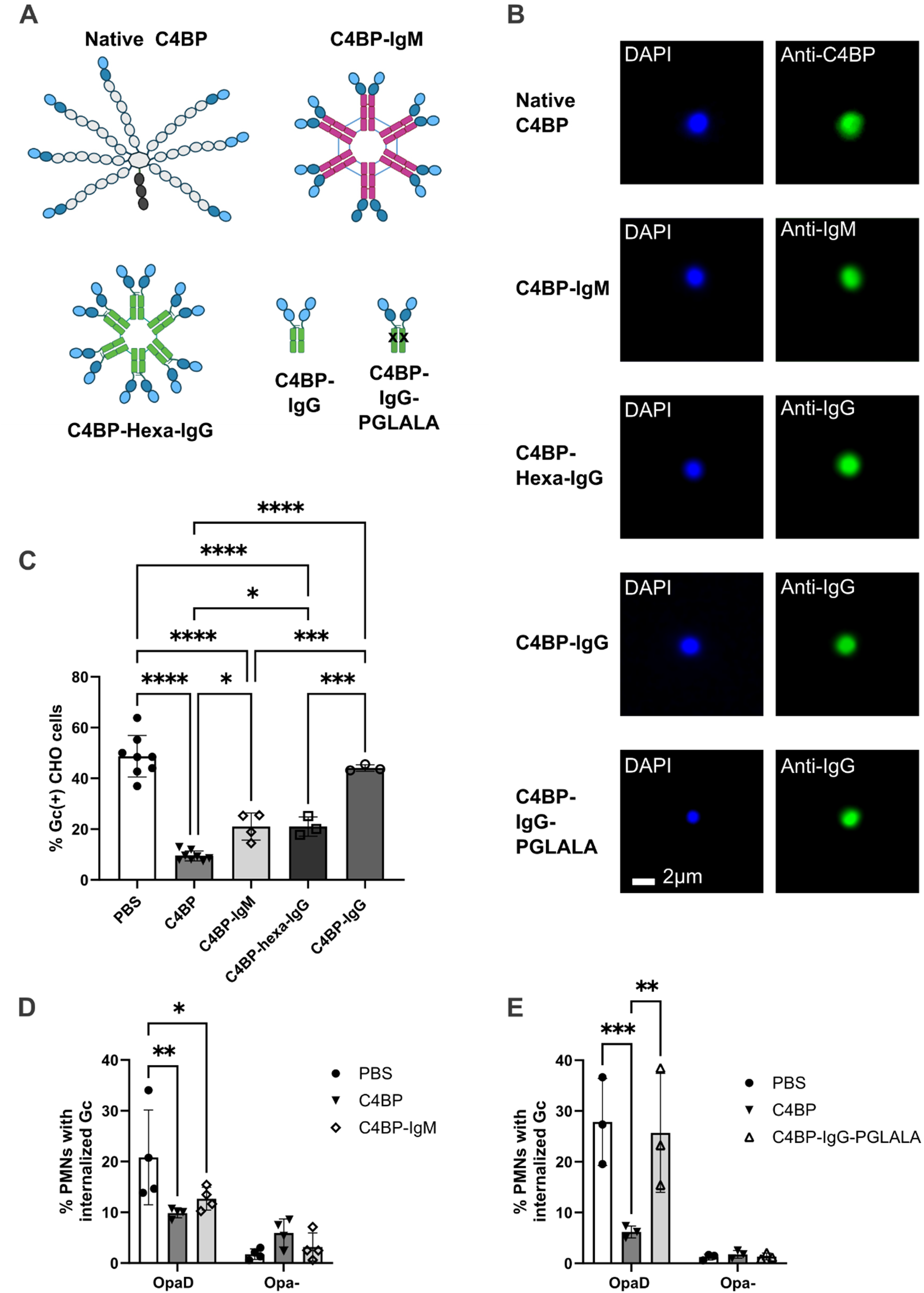
The size and multimeric nature of C4BP contribute to its ability to block interactions of Opa+ *N. gonorrhoeae* with CEACAM3 and phagocytosis by primary human neutrophils. (A) Structure of native C4BP and C4BP-based constructs. <u>Native C4BP</u>: blue = C4BPα CCP1-2 (CCP1 = light blue, CCP2 = dark blue); grey = C4BPα CCP3-8 and heptameric core; black = C4BPβ. <u>C4BP-IgM</u>: blue = C4BPα CCP1-2, pink = IgM Fc. <u>C4BP-Hexa-IgG</u>: blue = C4BPα CCP1-2; green = hexamerized IgG1 Fc. <u>C4BP-IgG</u>: blue = C4BPα CCP1-1; Green = IgG3 Fc. <u>C4BP-IgG-PGLALA</u>: blue = C4BPα CCP1-2; green = IgG1 Fc with L234A, L235A, and P329G Fc-silencing mutations, depicted by black x. (B) OpaD Gc was mixed with each of the indicated C4BP constructs at the following concentrations: PBS (vehicle), C4BP (100 µg/mL), C4BP-IgM (50 µg/mL), C4BP-Hexa-IgG (30 µg/mL), or C4BP-IgG (30 µg/mL) (C). Bacteria were fixed and processed for imaging flow cytometry, with DAPI to detect all bacteria and each construct detected with AlexaFluor 488-coupled antibodies as described in Materials and Methods. Scale bar = 2 µm. (C) CEACAM3-CHO cells were infected with TIV-labeled OpaD (MOI = 10 for each condition) that was pre-incubated with PBS (vehicle), C4BP, C4BP-IgM, C4BP-Hexa-IgG, or C4BP-IgG at the concentrations listed in B. Percent Gc-positive cells was calculated using imaging flow cytometry as in Figure 2. (D) Primary human neutrophils were infected with TIV-labeled OpaD or Opa-Gc (MOI = 1 for each condition) pre-incubated with PBS (vehicle), native C4BP, or C4BP-IgM, or with (E) PBS, native C4BP, or C4BP-IgG-PGLALA (100 µg/mL). Extracellular bacteria were detected using DyLight 650-coupled rabbit anti-Gc antibody without permeabilizing the cells. Internalization of Gc was assessed by imaging flow cytometry and quantified as the percent of neutrophils containing ≥ 1 intracellular bacterium (TIV-only). Graphs depict the mean +/− SD. Statistical significance was determined in (C) by one-way ANOVA with Tukey’s multiple comparisons and in (D-E) by paired two-way ANOVA with Tukey’s multiple comparisons, with comparisons made within each bacterial strain. *p<0.05, **p<0.01, ***p<0.001, ****p<0.0001. Non-significant comparisons are not shown.

We questioned if the same properties of C4BP that were important for blocking Opa-CEACAM3 interactions in transfected CHO cells were also important for inhibiting non-opsonic phagocytosis by primary human neutrophils. Neutrophils do not express FcµR, the receptor for IgM (Kubagawa et al., 2009). Therefore, it was possible to test if C4BP-IgM phenocopied the effects of native C4BP on non-opsonic phagocytosis in primary human neutrophils. Adherent primary human neutrophils primed with IL-8 were infected with OpaD that was previously incubated with C4BP-IgM, native C4BP, or vehicle alone. C4BP-IgM blocked the association and internalization of OpaD by primary human neutrophils similarly to native C4BP (Fig 7D).

Since neutrophils express FcγRs, it was not feasible to test the ability of the IgG-based C4BP constructs used above to inhibit bacterial association or internalization by primary human neutrophils. However, triple L234A/L235A/P329G Fc ‘silencing mutations (collectively called ‘PGLALA’) abrogate FcR engagement (Hale, 2024). Therefore, we generated C4BP-IgG1-PGLALA, in which the CCP1-2 regions of human C4BP replace the Fab region of IgG1, and the Fc region contains the PGLALA mutations. When preincubated with Gc, C4BP-IgG-PGLALA did not inhibit association or internalization of OpaD by neutrophils, while native C4BP did (Fig 7E). Importantly, none of the C4BP constructs used enhanced the phagocytosis of Opa-Gc, confirming their inability to serve as opsonins for phagocytic uptake (Fig 7D-E).

The ability of C4BP-IgM, but not C4BP-IgG-PGLALA, to block phagocytosis of OpaD implicates that the size and degree of multimerization of C4BP enable it to inhibit Opa-mediated phagocytosis by primary human neutrophils. Taken together with the results from CEACAM-transfected CHO cells, these results are consistent with a model in which C4BP sterically hinders Opa-CEACAM interaction, leading to enhanced survival of Opa+ Gc when confronted with human immune cell attack.

## Discussion

Gc binds the human complement inhibitor C4BP to evade soluble and cellular innate immunity, by preventing complement deposition and subsequent bactericidal lysis, and by blocking Opa-dependent, non-opsonic phagocytosis by neutrophils. To dissect the underlying mechanism, we generated CHO cells producing wild-type and variant human CEACAMs, and C4BP-based constructs of varying size and degree of multimerization. With this system, we found that C4BP inhibited the interaction between CEACAM-bearing CHO cells and Opa-expressing Gc, with a more pronounced effect on the granulocyte-restricted, phagocytic receptor CEACAM3 compared with the ubiquitously expressed ITIM-bearing CEACAM1. Using constructs that were engineered to have different numbers of extracellular IgC domains, C4BP had a greater inhibitory effect on the association of OpaD Gc with cells bearing receptors with shorter extracellular regions. We then engineered C4BP CCP1-2 based constructs that differ from native C4BP in their size and degree of multimerization, and found that the smaller, non-multimeric C4BP-IgG was unable to inhibit the interaction of OpaD Gc with CEACAM3-CHO cells or with primary human neutrophils. Together, these findings suggest that C4BP occludes the Opa-mediated interaction of Gc with CEACAM-bearing host cells, with a prominent effect on the neutrophil receptors CEACAM3 and CEACAM6, which enables the bacteria to evade phagocytic clearance.

For three different Opa-expressing clones of Gc, the inhibitory effect of C4BP was greater on CEACAM3 than CEACAM1, which was intriguing given that CEACAM3 is an activating, phagocytic receptor that acts through Syk kinase and Vav exchange factor for Rac GTPase (Sarantis & Gray-Owen, 2007; Schmitter et al., 2007), while CEACAM1 canonically elicits inhibitory signals via the SHP phosphatase (Boulton & Gray-Owen, 2002; Lee et al., 2008). We hypothesize that in intact neutrophils infected with C4BP-bound Gc, if CEACAM3 is inhibited more than CEACAM1, the signaling balance within the cell would be shifted towards a more inhibitory state. As a consequence, C4BP interference with Opa-CEACAM interaction on neutrophils would be expected to suppress downstream bactericidal activities such as ROS production, phagosome maturation, and cytokine release, which aligns with our prior findings (Werner et al., 2023). Moreover, CEACAM3 engagement drives an NF-κB-mediated transcriptional response that results in global cellular changes (Sintsova et al., 2014). Interestingly, there is evidence in heterologous overexpression systems that CEACAM1 engagement by Opa can also transactivate CEACAM3 (Sarantis & Gray-Owen, 2012). Thus, impeding both receptors is predicted to dampen neutrophil response to Gc. This would both enhance Gc survival and modulate the inflammatory milieu, which would support ongoing Gc infection. Future studies will examine how C4BP modulates CEACAM-derived neutrophil signaling and transcriptional responses to infection.

We found that CEACAM extracellular length is the main driver of the differences in C4BP-mediated inhibition between CEACAM1 and CEACAM3; there was no contribution of the ITAM region of CEACAM3 to the inhibitory effect of C4BP on Gc association with CHO cells. Replacing the IgV domain of CEACAM1 with that of CEACAM3 did not affect the inhibitory activity of CEACAM3, suggesting that Opa-binding sequences in the IgV domain do not contribute to the inhibitory effect of C4BP. CEACAM extracellular length, however, did contribute to the inhibitory effect of C4BP: we found that lengthening CEACAM3 decreased C4BP-mediated inhibition, and shortening CEACAM1 increased C4BP-mediated inhibition. Although these experiments were conducted in the heterologous CHO system, it points to extracellular interactions, rather than intracellular signaling, as the target of C4BP-mediated inhibition. Intriguingly, of the CEACAMs examined, C4BP most potently inhibited Opa interactions with GPI-anchored CEACAM6. This suggests that membrane dynamics and fluidity are also important contributors to the inhibitory effect of C4BP on Gc engagement of CEACAMs.

A difference between CEACAM1 and CEACAM3 that we did not explore is the ability of CEACAM1 to dimerize on the surface of the same cell and with neighboring cells, while CEACAM3 cannot (Bonsor et al., 2018; Gray-Owen & Blumberg, 2006). Transmembrane and IgV domains of CEACAM1 have both been implicated in dimerization (Klaile et al., 2009; Patel et al., 2013), though exact residues have not been fully defined. Reflective of this, we were unable to generate cells expressing a CEACAM3-based construct whose IgV domain was replaced with that of CEACAM1; the transfectants likely died because of perpetual dimerization and subsequent ITAM activation. If receptor dimerization contributes to the ability of CEACAM1 to bypass C4BP-mediated occlusion, differences in the ability of CEACAM1 and CEACAM3 to dimerize could explain why shortening CEACAM1 (CEACAM1S) did not fully recapitulate the inhibitory phenotype on CEACAM3. Generation of CEACAM1 and CEACAM1S mutants whose ability to dimerize is disrupted (without compromising the ability to bind Opa) could provide insight into the effect of CEACAM dimerization on C4BP-mediated inhibition.

C4BP is a large, branching protein whose most common isoform consists of 7 alpha chains and one beta chain. We found that the size and degree of multimerization of C4BP are important for its inhibitory effect on interactions of Opa with CEACAMs. CCP1-2 fused to multimeric Ig backbones (C4BP-IgM and C4BP-Hexa-IgG) were sufficient to recapitulate the effect of native C4BP on Opa-CEACAM3 interactions with CHO cells, as well as phagocytosis by primary human neutrophils. However, smaller dimeric C4BP-IgG constructs did not inhibit the interaction of Opa+ Gc with CEACAM3-CHO cells or primary human neutrophils.

Each of C4BP’s seven alpha chains contains binding sites for porin on CCP1, and porin is the most abundant protein on Gc’s outer membrane (Jarva et al., 2007; Johnston et al., 1976; Ram et al., 2001). Thus, interactions of each C4BP heptamer with the surface of Gc are multivalent in nature. Engineered C4BP alpha chain monomers do not bind efficiently to Gc, suggesting that the affinity of any one CCP for PorB is low, but avidity is amplified by the multimer (Ram et al., 2001; Werner et al., 2023). Each C4BP alpha chain is approximately 33 nm long and highly flexible (Dahlback et al., 1983; Kadava et al., 2025), which would allow the hexameric core of C4BP to be positioned high above the surface of Gc, and compress as cell membranes are pulled together via receptor-ligand interactions. In contrast to native C4BP, IgM and IgG Fc domains are more rigid in structure (Calvert et al., 2024; Remesh et al., 2018). Since C4BP-IgM and -Hexa-IgG were also able to block OpaD interactions with CEACAM3-CHO cells, it is unlikely that C4BP’s arm flexibility drives its inhibitory activity. We hypothesize that shortening the arms of C4BP would decrease its inhibitory activity, as it would present less of a physical barrier for host and bacterial cell membranes to come together and for Opa-CEACAM interactions to occur. The minimum degree of multimerization required to block Opa-CEACAM interactions also remains to be elucidated and will require attaching CCP domains 1 and 2 to scaffolds with different valencies.

C4BP-Ig fusions have been proposed as promising host-targeted therapeutics to combat Gc infection (Bettoni et al., 2019). In particular, the C4BP-IgM fusion protein has been shown to outcompete native C4BP for porin binding, drive complement-mediated killing of Gc, and increase susceptibility to antibiotics (Bettoni et al., 2021; Bettoni et al., 2019). Given that we showed C4BP-IgM decreases Gc phagocytosis by human neutrophils *ex vivo*, we hypothesize that as a therapeutic, C4BP-IgM may additionally dampen Opa-mediated neutrophilic inflammation. In the context of serum, this would likely be in balance with increased CR3-mediated phagocytosis and C5a-driven inflammatory programs, since increased C4BP-IgM drives C3 deposition on the surface of Gc and terminal complement activation (Bettoni et al., 2019). Enhancing CR3-mediated phagocytosis may not in fact lead to greater clearance of Gc, however, since we reported that Gc co-opts this receptor to enable it to be internalized without triggering strong neutrophil activation and bactericidal activities (Smirnov et al., 2023). All these potential effects of C4BP-IgM on Gc *in vivo* will be important to consider as its therapeutic potential is further evaluated and refined.

C4BP acquisition by Gc provides a powerful defense against complement and neutrophils. In this study, we show that C4BP occludes non-opsonic interactions between Gc Opa proteins and neutrophil CEACAMs. We propose a model in which C4BP most potently impedes interactions with receptors of shorter extracellular length, as the bacterial and neutrophil membranes would need to come together more closely for these receptor-ligand interactions to occur. These mechanisms can be used to inform C4BP-based therapeutics currently in development, as C4BP-based constructs of similar size and shape of native C4BP recapitulated the effect of C4BP on OpaD-CEACAM3 interactions. We predict that this C4BP-mediated inhibition of CEACAM binding dampens activating signaling, rewires transcriptional programs, and reduces the bactericidal capacity of neutrophils to ultimately protect Gc from neutrophil-mediated killing. Defining the role of C4BP in how Gc modulates host immune responses will enhance our understanding of the success of Gc within its obligate human host and reveal potential pathways to employ C4BP-based therapeutics for treating drug-resistant gonorrhea.

## Materials and Methods

### Bacterial strains and culture

*Neisseria gonorrhoeae* strains used in this study are in the FA1090 1-81-S2 Opa-negative (Opa-) background, which has all 11 *opa* genes chromosomally deleted (Ball & Criss, 2013). OpaD (Ball and Criss 2013), Opa60 (Martin et al., 2016), and OpaI (Alcott et al., 2022) contain single *opa* genes at the native *opaD* locus that are constitutively expressed due to incorporation of a non-phase variable signal sequence.

Bacterial lawns derived from individual colonies were cultured overnight at 37°C, 5% CO_2_ on Gonococcal Base Medium (GCB, Difco) agar plates containing Kellogg’s supplements 1 and 2 (Kellogg et al., 1963). Bacteria were grown to mid-log phase in Gonococcal base liquid medium (GCBL) also containing Kellogg’s supplements 1 and 2 with rotation at 37°C, with one back dilution and enrichment for piliated bacteria by selecting sedimented Gc as previously described (Ragland & Criss, 2019).

### Generation of Stable CEACAM-CHO cell lines

All CEACAM-encoding mammalian expression plasmids were synthesized by VectorBuilder and purified from *E. coli* stocks. CEACAM1, CEACAM3, or Control-CHOs were transfected using vectors encoding the cDNA from full-length human CEACAM1 (VB900000-5687rxc) or human CEACAM3 (VB900000-7471zkb) under a CAG promoter, or an empty control (VB010000-9288rhy). Each plasmid contains a dual EGFP/puromycin selection cassette. Mutant and chimeric CEACAM constructs sequences were synthesized on the same plasmid backbone. Sequences and descriptions of all CEACAM constructs are listed in Table S1.

Adherent PgsD-677 CHO K1 cells were acquired from ATCC (CRL-2244) and maintained in F12K medium (Gibco) supplemented with 10% heat-inactivated fetal bovine serum (HI-FBS) and antibiotics (1x antibiotic-antimycotic, Gibco) at 37°C, 5% CO_2_. Cells were transfected with plasmid DNA using Lipofectamine 3000 (Invitrogen) according to manufacturer’s protocol. Two days post-transfection, stable transfectants were selected using 10 µg/mL puromycin (Gibco) in regular culture media. Puromycin selection was maintained throughout culture.

### Sorting for CEACAM-expressing cells

Cells were lifted from tissue culture flasks by treatment with 5 mM ethylenediaminetetraacetic acid (EDTA) for 10 minutes at 37°C, 5% CO_2_ followed by gentle scraping with a plastic cell scraper. Cells were stained with mouse anti-Pan-CEACAM antibody (D14HD11, Abcam) for 30 minutes, washed, then stained with anti-mouse IgG-AlexaFluor (AF) 555 in culture medium. Cells that were positive for the vector uptake marker (EGFP) and the CEACAM expression marker (AlexaFluor (AF)555) were selected by flow cytometry cell sorting using Cytek Aurora or Sony MA900 cell sorters. Cells were re-plated and maintained in F12K medium supplemented with 10% HI-FBS, 1x antibiotic-antimycotic, and 10 µg/mL puromycin. Cells were monitored for CEACAM expression by immunofluorescence microscopy after sorting, after each freeze-thaw, and approximately once every 15 passages. All experiments were conducted using cells less than 30 passages post-transfection.

### Validation of CEACAM expression by immunofluorescence microscopy

Cells were seeded on tissue culture-treated plastic 2 days prior to staining. Cells were fixed in 4% paraformaldehyde, washed 3x in PBS, then blocked in PBS + 10% normal goat serum (NGS) (Gibco). Cells were stained with mouse anti-Pan-CEACAM for 30 minutes, washed 3x in PBS, then stained for 30 minutes with anti-mouse-IgG-AF555 and DAPI (1 µg/mL) counterstain. Images were acquired using a Keyence BZ-X series microscope using a 40x objective with DAPI and TRITC filter cubes.

### Generation and purification of C4BP-based constructs

Native C4BP from human serum (Complement Technologies #A109) was used in all experiments. C4BP-IgM was generated and purified as previously described (Bettoni et al., 2019). C4BP-Hexa-IgG and C4BP-IgG were commercially synthesized. To generate C4BP-Hexa-IgG, DNA sequences coding for CCP1-2 domains of C4BP and human IgG1-Fc were cloned in frame with N-terminal signal peptide sequence and C-terminal IgM tail piece into the viral based expression vector (pJL TRBO) (Bettoni et al., 2019; Lindbo, 2007; Yuan et al., 2022). DNA fragments were ordered from Thermo Fisher with GeneArt codon optimization for *Nicotiana benthamiana*. *N. benthamiana* variety deficient in xylosyl- and fucosyltransferase was vacuum infiltrated with agrobacterium strain carrying gene of interest (C4BP hexamer) into whole plants (Swope et al., 2021). Seven days post-infiltration, leaf material was harvested, homogenized, and filtered using miracloth (Millipore). C4BP-hexa-IgG was purified via protein A chromatography, then Capto core 400 chromatography was used to separate hexamer and monomer forms. Size and purity were verified by SDS-PAGE. C4BP-IgG was provided by Dr. Keith Wycoff (Planet Biotechnology, Inc.) and was similarly produced in *N. benthamiana* using previously described methods (Shaughnessy et al., 2022). C4BP-IgG-PGLALA was newly generated for this study. The nucleotide sequences encoding (from 5’ to 3’) human C4BP CCP domains 1 and 2 fused to the IgG1 hinge constant portion of human IgG1 with L234A/L235A/P329G mutations were synthesized and cloned into mammalian expression vector pcDNA 3.4 by GenScript (Piscataway, NJ). Expi-CHO cells (ThermoFisher) were transfected with the plasmid and the fusion protein, called C4BP-IgG-PGLALA, was purified from tissue culture supernatants using affinity chromatography over Protein A/G agarose. Purity and size of the obtained proteins were verified by SDS-PAGE. Protein concentration was quantified by absorption at 280 nm.

### Titrations of C4BP and C4BP-based constructs

Gc was incubated with the indicated concentrations of native C4BP, C4BP-IgM, C4BP-IgG, C4BP-Hexa-IgG, C4BP-IgG-PGLALA, or vehicle control in DPBS + 0.9 mM Ca + 0.49 mM Mg (“DPBS+/+”, Gibco 14040133) at 37°C for 20 minutes, incubated with fluorescent antibodies, and fixed in 1% PFA + 4 µg/mL DAPI counterstain. Native C4BP-treated conditions were stained with rabbit anti-C4BP (Novus Biologicals 88262) primary, followed by goat anti-rabbit AF488 (Invitrogen A-11008). C4BP-IgM-treated conditions were stained with anti-human-IgM-AF488 (Jackson 109-545-043), C4BP-IgG and hexa-IgG conditions were stained with anti-human IgG-AF488 (Invitrogen A-11013), and C4BP-IgG-PGLALA conditions were stained with Fcγ-specific anti-human IgG-AF488 (Jackson 109-545-008). Fluorescence intensity on DAPI+ bacteria was quantified by imaging flow cytometry.

### Bacterial incubation with C4BP or C4BP-based constructs and fluorescent labeling for infection

Gc (1 × 10^8^ CFU) were washed with DPBS+/+, then incubated in 250 µL DPBS+/+ containing 100 μg/mL native C4BP, 50 µg/mL C4BP-IgM, 30 μg/mL C4BP-IgG, 30 µg/mL C4BP-Hexa IgG, or vehicle alone for 20 minutes at 37°C. Bacteria were then labeled with Tag-it Violet^TM^ cell tracking and proliferation dye (TIV) (Biolegend) in DPBS+/+ for 15 minutes at 37°C. Bacteria were washed once more, then resuspended in the indicated infection medium (F12K + 10% FBS for CHO experiments, RPMI + 10% FBS for neutrophil experiments).

### Infection and imaging flow cytometry for Gc binding to CHO cells

Cells were seeded in a tissue culture treated 6-well plate 48 h pre-infection to yield a confluent monolayer containing approximately 1×10^6^ cells per well on the day of infection. 24 h prior to infection, cells were washed in DPBS (Gibco 14190144), and the selection/maintenance media was replaced with 2 mL antibiotic free infection medium (F12K + 10% FBS). 2 h prior to infection, medium was replaced with 1 mL fresh infection media.

CHO cells were synchronously infected with TIV-labeled Gc (prepared as described above) at the indicated MOIs via centrifugation (600xg, 4 minutes, 12°C). Infected cells were incubated at 37°C, 5% CO_2_ for 45 minutes. Cells were treated with 5 mM EDTA in DPBS for 10 minutes at 37°C, 5% CO_2_, then fixed by adding paraformaldehyde to 2% final concentration for 20 minutes. Cells were lifted using a rubber cell scraper, passed through a 70 µM nylon mesh cell strainer, then washed twice in PBS. Cells were stored at 4°C in the dark for less than three days before performing imaging flow cytometry. In experiments with unsorted cells where CEACAM staining was included, cells were blocked in PBS + 10% NGS, then stained with mouse anti-Pan-CEACAM primary antibody (D14HD11, Abcam), followed by goat anti-mouse-AF555. Data were collected using the ImageStream X MK II imaging flow cytometer.

CHO imaging flow data were analyzed with IDEAS v. 6.2 software. Single cells were gated in the brightfield channel by area and aspect ratio, and focused cells were selected using a gradient root mean square cutoff of ≥50. >3500 focused singlets were analyzed per experiment per condition. Single stained controls were used for fluorescence compensation and setting gates for CEACAM+ and TIV+ populations by fluorescence intensity.

Bacterial association with sorted cells was quantified using a spot counting-based approach. Bacterial particles associated with focused singlets were identified using mask definition Range(peak(Spot(M07, TIV, Bright, 1, 10, 1), TIV, Bright, 2.5). 4-80). Masking and spot counting functions were validated with random test images from each dataset. % Gc positive CHO cells is defined as the percent of total CHO cells containing one or more bacteria.

### Primary human neutrophil isolation

Venous blood from healthy human donors was collected in accordance with a protocol approved by the University of Virginia Institutional Review Board for Health Science Research (protocol #13909). Neutrophils were isolated as previously described (Ragland & Criss, 2019), using dextran sedimentation, Ficoll gradient centrifugation, and red blood cell lysis.

### Imaging flow cytometry for internalization of Gc by primary human neutrophils

Neutrophils were infected with TIV-labeled Gc, and bacterial internalization was assessed by imaging flow cytometry as previously described (Alcott et al., 2022; Smirnov et al., 2020; Werner et al., 2023). Extracellular Gc was identified via staining with rabbit anti-Gc antibody (Meridian B65111R) coupled to DyLight (DL) 650 (Thermo) in-house. Intracellular bacteria were identified as TIV(+) DL650(-) particles associated with focused, single neutrophils, and results are expressed as the percentage of neutrophils containing at least one intracellular Gc.

## Acknowledgements

We thank all members of the Criss Lab (UVA) for their feedback and advice, especially Ellie Martens for her assistance in validating spot-counting algorithms with test images. We thank M.W.B.’s thesis advisory committee for their guidance and expertise: Dr. Young Hahn, Dr. Sanja Arandjelovic, Dr. Dave Kashatus, and Dr. Ilya Leventhal (all UVA). Data for this manuscript were generated in the University of Virginia Flow Cytometry Core Facility (RRID:SCR_017829), which is partially supported by NIH P30CA044579. We thank the UVA Flow Cytometry Core staff, specifically Taylor Harper and Senthil Senguttuvan, for their assistance with cell sorting. Sony MA900 and the Cytek Aurora cell sorters were funded through the NIH instrument support program under 1S10OD028518 and 1S10OD034355, respectively. We thank Dr. Keith Wycoff (Planet Biotechnology, Inc.) for generating and providing C4BP-IgG, Dr. Kevin Whaley (Zabbio) and Dr. Jintang Du (K-Bio) for generating and providing C4BP-Hexa-IgG and Dr. Serena Bettoni for expressing and purifying C4BP-IgM. This work was supported by NIH R01A1097312 (AKC). M.W.B. was supported in part by NIH F31AI188753, NIH T32AI007046, and UVA Wagner Fellowship. A.J.C. was supported in part by NIH F31AI157528. AB is supported by grant from Swedish Research Council, 2022-00532). JS is supported by NIH R41 AI179576. SR is supported by NIH R43 AI186979. Research reported in this publication was supported by the National Institute of Allergy and Infectious Diseases of the National Institutes of Health. The content is solely the responsibility of the authors and does not necessarily represent the official views of the National Institutes of Health. Schematics *Created in BioRender*. Broden, M. (2026) https://BioRender.com/u3zn4io.

