## Supplemental materials for "C4BP occludes the non-opsonic interaction of *Neisseria gonorrhoeae* with human neutrophil CEACAMs"

**Supplemental Table 1: Sequences of all CEACAM constructs introduced into CHO cells by stable transfection**

Wild-type CEACAMs

| | NCBI RefSeq Number |
|---|---|
| **CEACAM1** | NM_001712.5 |
| **CEACAM3** | NM_001815.5 |
| **CEACAM6** | NM_002483.7 |

CEACAM constructs

| Construct/ Description | Sequence |
|---|---|
| **CEACAM3YYFF**<br><br>CEACAM3 with Y230F and Y241F mutations | ATGGGGCCCCCCTCAGCCTCTCCCCACAGAGAATGCATCCCCTGGCAGGGGCTTCTG<br>CTCACAGCCTCACTTCTAAACTTCTGGAACCCGCCCACCACTGCCAAGCTCACTATTG<br>AATCCATGCCGCTCAGTGTCGCAGAGGGGAAGGAGGTGCTTCTACTTGTCCACAATCT<br>GCCCCAGCATCTTTTTGGCTACAGCTGGTACAAAGGGGAAAGAGTGGATGGCAACAG<br>TCTAATTGTAGGATATGTAATAGGAACTCAACAAGCTACCCCAGGGGCCGCATACAGCG<br>GTCGAGAGACAATATACACCAATGCATCCCTGCTGATCCAGAATGTCACCCAGAATGAC<br>ATAGGATTCTACACCCTACAAGTCATAAAGTCAGATCTTGTGAATGAAGAAGCAACTGG<br>ACAGTTCCATGTATACCAAGAAAATGCCCCAGGCCTTCCTGTGGGGGCCGTCGCCGG<br>CATCGTGACCGGGGTCCTGGTCGGAGTGGCGCTGGTGGCCGCGCTGGTGTGTTTCC<br>TGCTCCTTGCCAAAACTGGAAGAACCAGCATCCAGCGTGACCTCAAGGAGCAGCAGC<br>CCCAAGCCCTTGCCCCTGGCCGTGGTCCCTCCCACAGCTCTGCCTTCTCGATGTCCC<br>CTCTCTCCACTGCCCAGGCCCCCCTACCCAACCCCAGGACAGCAGCTTCCATCTTTG<br>AGGAATTGCTAAAACATGACACAAACATTTTCTGCCGGATGGACCACAAAGCAGAAGT<br>GGCTTCTTAG |
| **CEACAM1-3N**<br><br><br>CEACAM3 Signal sequence and IgV domain (codons 1-142), CEACAM1 IgC, Transmembrane, and ITIM domains (143-527) | ATGGGGCCCCCCTCAGCCTCTCCCCACAGAGAATGCATCCCCTGGCAGGGGCTTCTG<br>CTCACAGCCTCACTTCTAAACTTCTGGAACCCGCCCACCACTGCCAAGCTCACTATTG<br>AATCCATGCCGCTCAGTGTCGCAGAGGGGAAGGAGGTGCTTCTACTTGTCCACAATCT<br>GCCCCAGCATCTTTTTGGCTACAGCTGGTACAAAGGGGAAAGAGTGGATGGCAACAG<br>TCTAATTGTAGGATATGTAATAGGAACTCAACAAGCTACCCCAGGGGCCGCATACAGCG<br>GTCGAGAGACAATATACACCAATGCATCCCTGCTGATCCAGAATGTCACCCAGAATGAC<br>ATAGGATTCTACACCCTACAAGTCATAAAGTCAGATCTTGTGAATGAAGAAGCAACTGG<br>ACAGTTCCATGTATACCAAGAGCTGCCCAAGCCCTCCATCTCCAGCAACAACTCCAAC<br>CCTGTGGAGGACAAGGATGCTGTGGCCTTCACCTGTGAACCTGAGACTCAGGACACA<br>ACCTACCTGTGGTGGATAAACAATCAGAGCCTCCCGGTCAGTCCCAGGCTGCAGCTG<br>TCCAATGGCAACAGGACCCTCACTCTACTCAGTGTCACAAGGAATGACACAGGACCCT<br>ATGAGTGTGAAATACAGAACCCAGTGAGTGCGAACCGCAGTGACCCAGTCACCTTGA<br>ATGTCACCTATGGCCCGGACACCCCCACCATTTCCCCTTCAGACACCTATTACCGTCC<br>AGGGGCAAACCTCAGCCTCTCCTGCTATGCAGCCTCTAACCCACCTGCACAGTACTCC<br>TGGCTTATCAATGGAACATTCCAGCAAAGCACACAAGAGCTCTTTATCCCTAACATCAC<br>TGTGAATAATAGTGGATCCTATACCTGCCACGCCAATAACTCAGTCACTGGCTGCAACA<br>GGACCACAGTCAAGACGATCATAGTCACTGAGCTAAGTCCAGTAGTAGCAAAGCCCCA<br>AATCAAAGCCAGCAAGACCACAGTCACAGGAGATAAGGACTCTGTGAACCTGACCTG<br>CTCCACAAATGACACTGGAATCTCCATCCGTTGGTTCTTCAAAAACCAGAGTCTCCCG<br>TCCTCGGAGAGGATGAAGCTGTCCCAGGGCAACACCACCCTCAGCATAAACCCTGTC<br>AAGAGGGAGGATGCTGGGACGTATTGGTGTGAGGTCTTCAACCCAATCAGTAAGAAC<br>CAAAGCGACCCCATCATGCTGAACGTAAACTATAATGCTCTACCACAAGAAAATGGCCT<br>CTCACCTGGGGCCATTGCTGGCATTGTGATTGGAGTAGTGGCCCTGGTTGCTCTGATA<br>GCAGTAGCCCTGGCATGTTTTCTGCATTTCGGGAAGACCGGCAGGGCAAGCGACCAG<br>CGTGATCTCACAGAGCACAAACCCTCAGTCTCCAACCACACTCAGGACCACTCCAATG<br>ACCCACCTAACAAGATGAATGAAGTTACTTATTCTACCCTGAACTTTGAAGCCCAGCAA<br>CCCACACAACCAACTTCAGCCTCCCCATCCCTAACAGCCACAGAAATAATTTATTCAGA<br>AGTAAAAAAGCAGTAA |
| **CEACAM3L** | ATGGGGCCCCCCTCAGCCTCTCCCCACAGAGAATGCATCCCCTGGCAGGGGCTTCTG<br>CTCACAGCCTCACTTCTAAACTTCTGGAACCCGCCCACCACTGCCAAGCTCACTATTG |

| | |
|---|---|
| CEACAM3 Signal sequence and IgV domain (1-142), CEACAM1 IgC1-3 Domains (143-428), CEACAM3 Transmembrane and ITAM domains (429-526) | AATCCATGCCGCTCAGTGTCGCAGAGGGGAAGGAGGTGCTTCTACTTGTCCACAATCT<br>GCCCCAGCATCTTTTTGGCTACAGCTGGTACAAAGGGGAAAGAGTGGATGGCAACAG<br>TCTAATTGTAGGATATGTAATAGGAACTCAACAAGCTACCCCAGGGGCCGCATACAGCG<br>GTCGAGAGACAATATACACCAATGCATCCCTGCTGATCCAGAATGTCACCCAGAATGAC<br>ATAGGATTCTACACCCTACAAGTCATAAAGTCAGATCTTGTGAATGAAGAAGCAACTGG<br>ACAGTTCCATGTATACCAAGAGCTGCCCAAGCCCTCCATCTCCAGCAACAACTCCAAC<br>CCTGTGGAGGACAAGGATGCTGTGGCCTTCACCTGTGAACCTGAGACTCAGGACACA<br>ACCTACCTGTGGTGGATAAACAATCAGAGCCTCCCGGTCAGTCCCAGGCTGCAGCTG<br>TCCAATGGCAACAGGACCCTCACTCTACTCAGTGTCACAAGGAATGACACAGGACCCT<br>ATGAGTGTGAAATACAGAACCCAGTGAGTGCGAACCGCAGTGACCCAGTCACCTTGA<br>ATGTCACCTATGGCCCGGACACCCCCACCATTTCCCCTTCAGACACCTATTACCGTCC<br>AGGGGCAAACCTCAGCCTCTCCTGCTATGCAGCCTCTAACCCACCTGCACAGTACTCC<br>TGGCTTATCAATGGAACATTCCAGCAAAGCACACAAGAGCTCTTTATCCCTAACATCAC<br>TGTGAATAATAGTGGATCCTATACCTGCCACGCCAATAACTCAGTCACTGGCTGCAACA<br>GGACCACAGTCAAGACGATCATAGTCACTGAGCTAAGTCCAGTAGTAGCAAAGCCCCA<br>AATCAAAGCCAGCAAGACCACAGTCACAGGAGATAAGGACTCTGTGAACCTGACCTG<br>CTCCACAAATGACACTGGAATCTCCATCCGTTGGTTCTTCAAAAACCAGAGTCTCCCG<br>TCCTCGGAGAGGATGAAGCTGTCCCAGGGCAACACCACCCTCAGCATAAACCCTGTC<br>AAGAGGGAGGATGCTGGGACGTATTGGTGTGAGGTCTTCAACCCAATCAGTAAGAAC<br>CAAAGCGACCCCATCATGCTGAACGTAAACTATAATGCTCTACCACAAGAAAATGGCCT<br>CTCACCTGGGATCGTGACCGGGGTCCTGGTCGGAGTGGCGCTGGTGGCCGCGCTG<br>GTGTGTTTCCTGCTCCTTGCCAAAACTGGAAGAACCAGCATCCAGCGTGACCTCAAG<br>GAGCAGCAGCCCCAAGCCCTTGCCCCTGGCCGTGGTCCCTCCCACAGCTCTGCCTT<br>CTCGATGTCCCCTCTCTCCACTGCCCAGGCCCCCCTACCCAACCCCAGGACAGCAGC<br>TTCCATCTATGAGGAATTGCTAAAACATGACACAAACATTTACTGCCGGATGGACCACA<br>AAGCAGAAGTGGCTTCTTAG |
| **CEACAM3XL**<br><br>CEACAM3 Signal Sequence and IgV domain (1-142),<br>EL Linker (143-144),<br>CEACAM5 IgC1-6 domains codon optimized for *C. griseus* (145-675),<br>CEACAM3 transmembrane and ITAM domains (676-786) | ATGGGGCCCCCCTCAGCCTCTCCCCACAGAGAATGCATCCCCTGGCAGGGGCTTCTG<br>CTCACAGCCTCACTTCTAAACTTCTGGAACCCGCCCACCACTGCCAAGCTCACTATTG<br>AATCCATGCCGCTCAGTGTCGCAGAGGGGAAGGAGGTGCTTCTACTTGTCCACAATCT<br>GCCCCAGCATCTTTTTGGCTACAGCTGGTACAAAGGGGAAAGAGTGGATGGCAACAG<br>TCTAATTGTAGGATATGTAATAGGAACTCAACAAGCTACCCCAGGGGCCGCATACAGCG<br>GTCGAGAGACAATATACACCAATGCATCCCTGCTGATCCAGAATGTCACCCAGAATGAC<br>ATAGGATTCTACACCCTACAAGTCATAAAGTCAGATCTTGTGAATGAAGAAGCAACTGG<br>ACAGTTCCATGTATACCAAGAGCTGCCCAAACCTTCTATCAGCTCTAACAACTCTAAGC<br>CTGTGGAGGATAAGGATGCCGTGGCTTTTACATGCGAGCCTGAGACCCAGGACGCCA<br>CCTACCTGTGGTGGGTGAACAACCAGAGCCTGCCCGTGAGCCCCCGCCTGCAGCTG<br>TCTAACGGAAATAGGACCCTGACCCTGTTCAATGTGACACGAAACGACACAGCCAGCT<br>ATAAGTGTGAGACCCAGAACCCAGTGAGCGCACGGAGATCAGATTCTGTGATCCTGAA<br>CGTGCTGTATGGGCCCGACGCCCCAACAATTTCCCCCCTGAACACCAGTTATCGCAG<br>CGGCGAGAACCTGAACCTGAGCTGCCACGCCGCTTCCAATCCCCCCGCCCAGTACTC<br>TTGGTTCGTGAATGGCACATTCCAGCAGAGCACCCAGGAGCTGTTCATTCCTAATATCA<br>CAGTGAATAATTCTGGCTCTTACACCTGTCAGGCTCACAACTCTGACACTGGCCTGAAT<br>AGGACTACAGTGACTACCATCACCGTGTACGCTGAACCACCTAAGCCCTTCATTACTAG<br>CAACAATTCTAATCCCGTGGAAGACGAGGATGCCGTGGCTCTGACATGCGAGCCCGA<br>GATCCAGAACACCACATACCTGTGGTGGGTGAACAACCAGTCCCTGCCCGTGTCTCC<br>AAGACTGCAGCTGTCTAACGACAACAGGACCCTGACCCTGCTGTCTGTGACTCGCAA<br>CGACGTGGGCCCATACGAGTGCGGAATTCAGAACGAGCTGTCCGTGGACCACTCTGA<br>TCCTGTGATCCTGAACGTGCTGTACGGCCCTGACGACCCTACTATTTCTCCCTCTTACA<br>CATACTACAGGCCCGGCGTGAACCTGTCCCTGAGTTGTCATGCCGCTAGTAATCCCCC<br>TGCCCAGTACAGCTGGCTGATCGATGGCAATATCCAGCAGCATACCCAGGAGCTGTTC<br>ATCTCCAATATCACAGAGAAGAACTCTGGCCTGTACACCTGCCAGGCCAACAATTCCG<br>CTTCTGGCCATTCTCGCACCACCGTGAAGACCATCACAGTGAGCGCCGAGCTTCCCA<br>AGCCCTCTATCAGCTCTAATAATTCCAAGCCCGTGGAAGACAAGGACGCCGTGGCTTT<br>CACCTGTGAGCCTGAGGCTCAGAACACAACCTACCTCTGGTGGGTGAATGGCCAGTC<br>TCTGCCCGTCAGCCCTAGGCTGCAGCTGTCCAATGGTAATCGGACCCTGACACTGTTC<br>AACGTCACCAGGAACGACGCCAGGGCTTACGTGTGCGGCATCCAGAACAGCGTGAG<br>CGCCAACCGGAGTGACCCTGTGACCCTGGACGTGCTGTATGGTCCTGACACCCCCAT<br>CATCAGCCCACCCGACTCCTCCTACCTCAGCGGGGCCAATCTGAACCTGTCTTGCCA<br>CTCTGCCTCTAATCCTTCCCCCCAGTACAGTTGGAGGATCAACGGAATCCCCCAGCAG<br>CACACCCAGGTGCTTTTCATTGCCAAGATCACCCCAAATAACAATGGCACATATGCCTG<br>CTTTGTGAGCAACCTGGCTACCGGCCGGAACAACTCCATCGTGAAGTCCATTACAGTG<br>TCTGAAAATGCCCCAGGCCTTCCTGTGGGGGCCGTCGCCGGCATCGTGACCGGGGT |

| | |
|---|---|
| | CCTGGTCGGAGTGGCGCTGGTGGCCGCGCTGGTGTGTTTCCTGCTCCTTGCCAAAA<br>CTGGAAGAACCAGCATCCAGCGTGACCTCAAGGAGCAGCAGCCCCAAGCCCTTGCC<br>CCTGGCCGTGGTCCCTCCCACAGCTCTGCCTTCTCGATGTCCCCTCTCTCCACTGCC<br>CAGGCCCCCCTACCCAACCCCAGGACAGCAGCTTCCATCTATGAGGAATTGCTAAAAC<br>ATGACACAAACATTTACTGCCGGATGGACCACAAAGCAGAAGTGGCTTCTTAG |
| **CEACAM1S**<br><br>CEACAM1 IgV domain (1-142), CEACAM1 Transmembrane and ITIM domains (143-256) | ATGGGGCACCTCTCAGCCCCACTTCACAGAGTGCGTGTACCCTGGCAGGGGCTTCTG<br>CTCACAGCCTCACTTCTAACCTTCTGGAACCCGCCCACCACTGCCCAGCTCACTACTG<br>AATCCATGCCATTCAATGTTGCAGAGGGGAAGGAGGTTCTTCTCCTTGTCCACAATCT<br>GCCCCAGCAACTTTTTGGCTACAGCTGGTACAAAGGGGAAAGAGTGGATGGCAACCG<br>TCAAATTGTAGGATATGCAATAGGAACTCAACAAGCTACCCCAGGGCCCGCAAACAGC<br>GGTCGAGAGACAATATACCCCAATGCATCCCTGCTGATCCAGAACGTCACCCAGAATG<br>ACACAGGATTCTACACCCTACAAGTCATAAAGTCAGATCTTGTGAATGAAGAAGCAACT<br>GGACAGTTCCATGTATACCCGGTAAACTATAATGCTCTACCACAAGAAAATGGCCTCTC<br>ACCTGGGGCCATTGCTGGCATTGTGATTGGAGTAGTGGCCCTGGTTGCTCTGATAGCA<br>GTAGCCCTGGCATGTTTTCTGCATTTCGGGAAGACCGGCAGGGCAAGCGACCAGCGT<br>GATCTCACAGAGCACAAACCCTCAGTCTCCAACCACACTCAGGACCACTCCAATGACC<br>CACCTAACAAGATGAATGAAGTTACTTATTCTACCCTGAACTTTGAAGCCCAGCAACCC<br>ACACAACCAACTTCAGCCTCCCCATCCCTAACAGCCACAGAAATAATTTATTCAGAAGT<br>AAAAAAGCAGTAA |

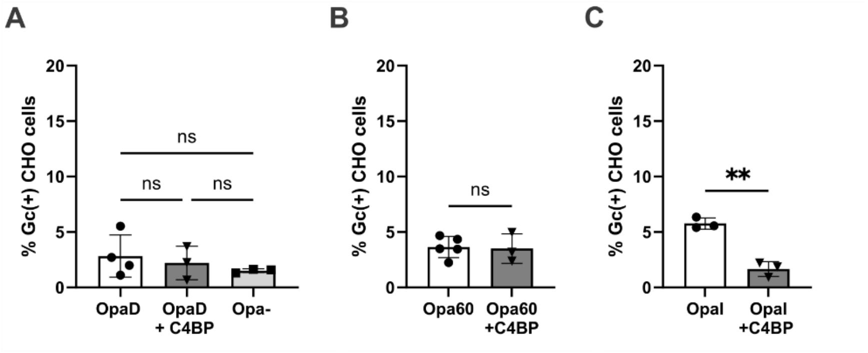


**Supplemental Figure 1: Control-CHO cells exhibit minimal interaction with Gc, regardless of bacterial binding to C4BP.**

CHO cells transfected with an empty vector (Control-CHO cells) were infected with TIV-labeled Gc, and the percent of cells positive for Gc was quantified from imaging flow cytometry images as in Figure 2. (A) Control-CHO cells infected with OpaD ± C4BP or Opa-, each at MOI = 10. (B) Control-CHO cells infected with Opa60 ± C4BP, each at MOI = 10. (C) Control-CHO cells infected with OpaI ± C4BP, each at MOI = 5. Graphs depict the mean +/- SD. Statistical significance was determined in (A) by one-way ANOVA with Tukey's multiple comparisons and in (B-C) by unpaired T-test. **p<0.01, ns = not significant.

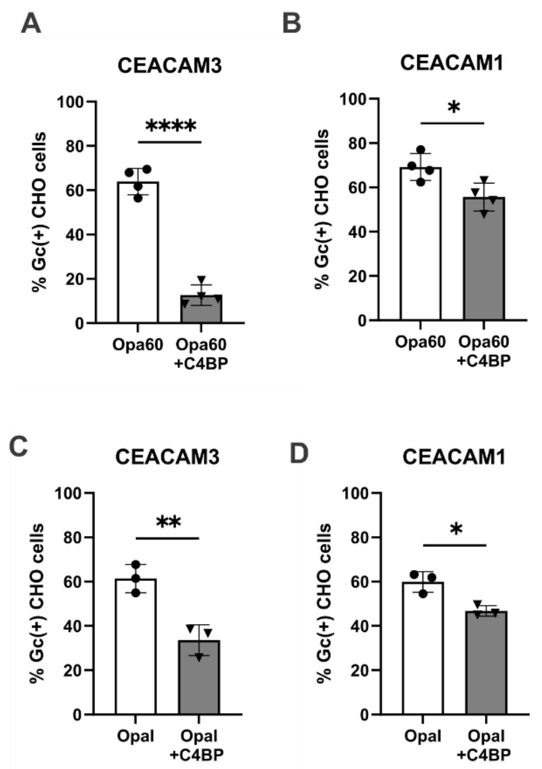


**Supplemental Figure 2: C4BP more potently inhibits interactions of Opa60 and OpaI with CEACAM3-CHO cells than CEACAM1-expressing cells.**

CEACAM1-CHO or CEACAM3-CHO cells were infected with TIV-labeled Gc, and bacterial binding was assessed using imaging flow cytometry. (A) CEACAM3-CHO cells were infected with Opa60 ± C4BP (MOI = 10 for each condition). (B) CEACAM1-CHO cells were infected with Opa60 ± C4BP (MOI = 5 for each condition). (C) CEACAM3-CHO cells were infected with OpaI ± C4BP (MOI = 5 for each condition). (D) CEACAM1-CHO cells were infected with OpaI ± C4BP. MOI = 5. (A-D) Percent Gc-positive cells were calculated using imaging flow cytometry as in Figure 2. Graphs depict the mean +/- SD. Statistical significance was determined by unpaired T-test. *$p<0.05$, **$p<0.01$, ****$p<0.0001$.

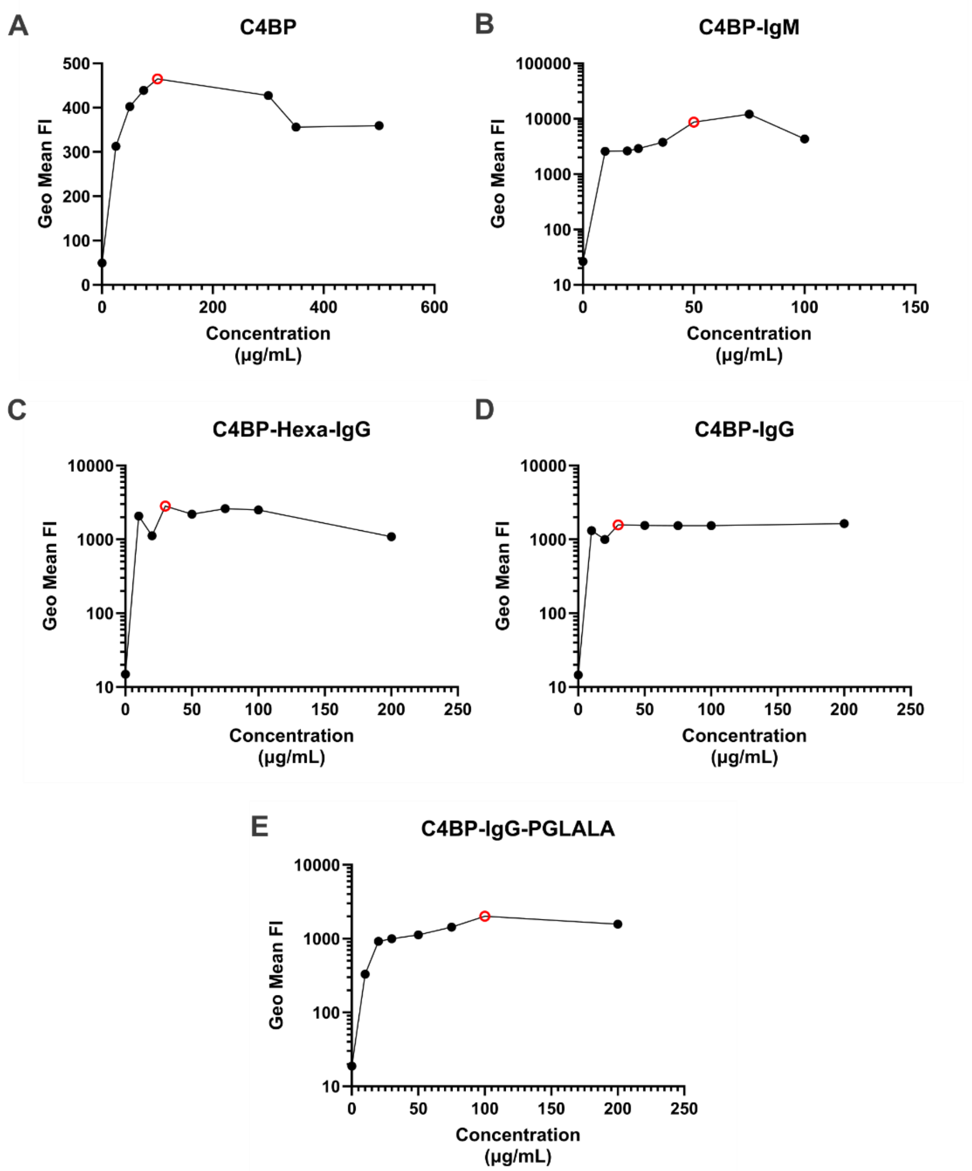


**Supplemental Figure 3: Titration of the binding of C4BP and related constructs to OpaD Gc.**

OpaD was incubated with the indicated concentrations of C4BP and related constructs at 37°C for 20 minutes, washed, then stained with the following: (A), C4BP: rabbit anti-C4BP primary, Anti-rabbit-AlexaFluor 488 (AF488) secondary. (B), C4BP-IgM: Anti-human IgM-AF488. (C),C4BP-Hexa-IgG: Anti-Human IgG-AF488. (D), C4BP-IgG: Anti-Human IgG-AF488. (E), C4BP-IgG-PGLALA: Anti-human IgG Fcγ-AF488. (A-E) Brightfield and DAPI counterstain were used to identify bacterial singlets by imaging flow cytometry. Data are presented as the geometric mean fluorescence intensity of AF488. The minimum concentration for maximum binding of each construct to Gc was used for the experiments in Figure 7, designated by the red open circle on each graph using that construct.
